# Cryobanking of human distal lung epithelial cells for preservation of their phenotypic and functional characteristics

**DOI:** 10.1101/2021.11.15.468402

**Authors:** Bindu Konda, Apoorva Mulay, Changfu Yao, Edo Israely, Stephen Beil, Carissa A. Huynh, Warren G. Tourtellotte, Reinaldo Rampolla, Peter Chen, Gianni Carraro, Barry R. Stripp

**Affiliations:** Lung Institute, Department of Medicine, Cedars-Sinai Medical Center, Los Angeles, CA, USA; Regenerative Medicine Institute, Cedars-Sinai Medical Center, Los Angeles, CA, USA; Department of Pathology, Cedars-Sinai Medical Center, Los Angeles, Ca, USA

**Keywords:** Epithelial progenitor cells, Organoids, Cryopreservation, single-cell RNA sequencing

## Abstract

The epithelium lining airspaces of the human lung is maintained by regional stem cells including basal cells of pseudostratified airways and alveolar type 2 pneumocytes (AT2) of the alveolar gas-exchange region. Despite effective methods for long-term preservation of airway basal cells, methods for efficient preservation of functional epithelial cell types of the distal gas-exchange region are lacking. Here we detail a method for cryobanking of epithelial cells from either mouse or human lung tissue for preservation of their phenotypic and functional characteristics. Flow cytometric profiling, epithelial organoid-forming efficiency, and single cell transcriptomic analysis, were used to compare cells recovered from cryopreserved tissue with those of freshly dissociated tissue. Alveolar type 2 cells within single cell suspensions of enzymatically digested cryobanked distal lung tissue retained expression of the pan-epithelial marker CD326 and the AT2 cell surface antigen recognized by monoclonal antibody HTII-280, allowing antibody-mediated enrichment and downstream analysis. Isolated AT2 cells from cryobanked tissue were comparable with those of freshly dissociated tissue both in their single cell transcriptome and their capacity for in vitro organoid formation in 3D cultures. We conclude that the cryobanking method described herein allows long-term preservation of distal human lung tissue for downstream analysis of lung cell function and molecular phenotype, and is ideally suited for creation of an easily accessible tissue resource for the research community.

## INTRODUCTION

Procurement of fresh lung tissue from both healthy and diseased donors is an integral part of research focused on human lung biology and disease. The expanding repertoire of advanced omics approaches allowing molecular phenotyping of lung cells at single-cell resolution[1-3], coupled with advances in disease modeling using organoid-based three-dimensional culture systems[4-7], has increased demand for this limited tissue resource. Comparisons made between lung cells isolated from either control tissue from donors lacking prior clinical evidence of lung disease, or specimens recovered from patients either through bronchoscopy, biopsy or following transplantation, have yielded novel insights into the cell biology and pathophysiology of the human lung [8-15]. Collection of sufficient biological replicates of fresh tissue is challenging and typically requires establishment of tissue procurement/sharing consortia[16, 17]. In addition, omics studies performed with high biological replicates at the time of tissue procurement results in technical batch effects that must be corrected post-hoc, with associated reductions in data quality and reliability [18]. Accordingly, there is an urgent need to validate efficient methodologies for tissue banking that preserve the molecular phenotype and function of lung cells within valuable donated tissue samples.

More applied applications of primary lung tissue samples include the development of epithelial culture models for optimization of either cell-or gene-based corrective therapies. Cell-based therapies have been adopted for diseases involving a variety of organs and cell types [19-31], and in combination with therapeutic gene correction of long-lived lung epithelial progenitors, may represent a plausible approach for monogenic lung diseases such as cystic fibrosis [32-34]. However, realization of stem cell-based therapeutic approaches for lung regenerative medicine will require development and validation of reproducible protocols allowing standardization and scalability, as has been demonstrated in other tissues [35-38].

Herein, we demonstrate a novel approach for human lung tissue processing and preservation that allows on-demand access to donor tissue samples. We show that tissue dissociation of Cryobanked mouse and human lung tissue samples yield epithelial cell types whose molecular phenotype and functional properties in organoid-based culture models are equivalent to those of freshly dissociated tissue. Application of this method to tissue samples provided by otherwise healthy lung tissue donors, or patients donating explanted tissue for research at the time of transplantation or partial surgical resection, has potential to significantly expand the accessibility of these precious samples for basic and translational research into human lung cell biology and disease.

## MATERIALS AND METHODS

### Animals

Experiments were performed with pathogen free C57Bl/6 mice in accordance with institutional IACUC approval.

### Human lung samples

Human lung tissue was obtained from deceased tissue donors in compliance with consent procedures developed by the International Institute for the Advancement of Medicine (IIAM) and approved by the Cedars-Sinai Medical Center Internal Review Board.

Detailed description of the materials and methods are available as an online supplement.

### Freezing distal regions of the lung

Lung tissue was diced into pieces of approximately 1 cm^3^ and placed in a 50 mL conical tube, washed to remove blood and epithelial lining fluid, and minced into pieces of approximately 3-4 millimeters in diameter. Between 1 and 1.5 grams of tissue was added to a 2 mL cryovial with 1 mL of CryoStor cryopresearvative media. Vials placed in a cell freezing container filled with isopropyl alcohol, were left at -80 °C overnight. The following day, vials were transferred to liquid nitrogen for long-term storage.

Note: The tissue needs to be completely submerged in freezing medium in order to prevent loss of cell viability during the freezing process.

### Enrichment and subsetting of epithelial progenitor cells from fresh or frozen lung tissue

Enrichment and fractionation of small airway and alveolar progenitor cells from distal lung tissue was performed using appropriate cell surface markers, as previously described [39]. Tissue was thawed by placing the vial at 37 °C for 1 min and transferred into a sterile 50 ml tube with 10ml of HBSS and centrifuged at 500 g for 5 min. Tissue was then transferred to a sterile petri dish, finely minced using single sided razor blade, and enzymatically digested using Liberase (0.05 mg/mL), DNase (0.025 mg/mL) for 40-60 min at 37 °C, to obtain a single cell suspension.

NOTE: Incubation time with the enzymes can vary depending on the type or condition of the tissue. For example, enzymatic digestion of normal tissue takes approximately 45 min. However, fibrotic tissue from Idiopathic pulmonary fibrosis samples can require a longer incubation time of up to 60 min. Therefore, monitor tissue carefully during this step.

Depletion of mesenchymal and immune fractions and enrichment of HT II-280+ cells, was performed as previously described[39, 40].

### Large scale organoid expansion

Organoids were cultured as previously described[39].

Organoids were passaged by washing once with PBS and Matrigel dissolved by the addition of Corning cell recovery solution. Cells were collected in FACS buffer (supplemental data Table 2) and centrifuged at 600 x g for 5 minutes at 4 °C, followed by staining for FACS. Primary antibodies were added at the required concentration to each 1 × 10^7^ cells in 1 ml of HBSS+ buffer and incubated for 30 minutes at 4 °C (Table 3). For cultures, 5 × 10^3^ enriched epithelial cells were added to a sterile 1.5 mL tube along with 7.5 × 10^4^ MRC-5 cells (human lung fibroblast cell line), cell were centrifuged at 500 x g for 5 minutes at 4 °C. NOTE: Cell counts determined by FACS must be confirmed by manual counting with a hemocytometer. Cell pellets are resuspended in antibiotic-supplemented media and placed on ice. Next, an equal volume of 1x growth factor depleted basement membrane matrix medium (Matrigel) was added. Samples should be kept on ice to avoid premature polymerization of Matrigel. The cell suspension is then transferred into a 24-well Transwell insert (0.4 μm pore-size; 1.4 × 10^4^ cells/cm^2^) (supplemental data Table 4), taking care not to introduce air bubbles. After incubation at 37 °C for 30-45 minutes, Transwell’s are placed in a culture plate containing 600 μL of pre-warmed growth medium.

NOTE: Antimycotic agents (1μg/ml) and Gentamicin (50μg/ml) should be included in culture media for the first 24 hours, and 10 μM Rho kinase inhibitor added for the first 72 hours after initial plating. Cultures should be kept at 37 °C in a 5% CO_2_ incubator for up to 30 days, during which time the media should be changed every 48 h.

### Isolation of cells from Mouse Lung

See supplemental methods.

### Single cell RNA sequencing

Single cells were captured from FACS-enriched epithelial fractions (CD45^**-**^, CD31^**-**^, CD326^**+**^) of fresh and frozen tissue using the 10x Chromium platform and used for the generation of scRNA-Seq libraries according to the manufacturer’s instructions (10X Genomics). Briefly, cell suspensions were loaded on a Chromium Controller instrument (10X Genomics) to generate single-cell Gel Bead-In-Emulsions (GEMs). Reverse transcription (RT) was performed in a Veriti 96-well thermal cycler. Barcoded sequencing libraries were quantified by quantitative PCR using the KAPA Library Quantification Kit (KAPA Biosystems, Wilmington, MA), and sequencing performed on a NovaSeq 6000 (Illumina). Cell Ranger software (10X Genomics) was used for mapping and barcode filtering. Briefly, the raw reads were aligned to the transcriptome using STAR(5), using a hg38 transcriptome reference from Ensemble 93 annotation. Expression counts for each gene in all samples were collapsed and normalized to unique molecular identifier (UMI) counts. Data analysis was performed with Conos, a tool developed for joint analysis of heterogeneous datasets.

## RESULTS

### Mouse lung Epcam^+^ cell fractionation comparison fresh tissue Vs frozen tissue Vs frozen cells

Using a newly devised experimental approach (Fig. 1A-E), we have optimized the extraction of cryopreserved epithelial progenitor cells from human and mouse lung. In this workflow, lung tissue is enzymatically digested using a combination of Liberase and Dispase. Immune (CD45^**+**^) and endothelial (CD31^**+**^) cells are then removed using magnetic beads. Finally, using specific epithelial surface markers, cells undergo FACS isolation, before being processed for downstream applications.

**Figure 1:**
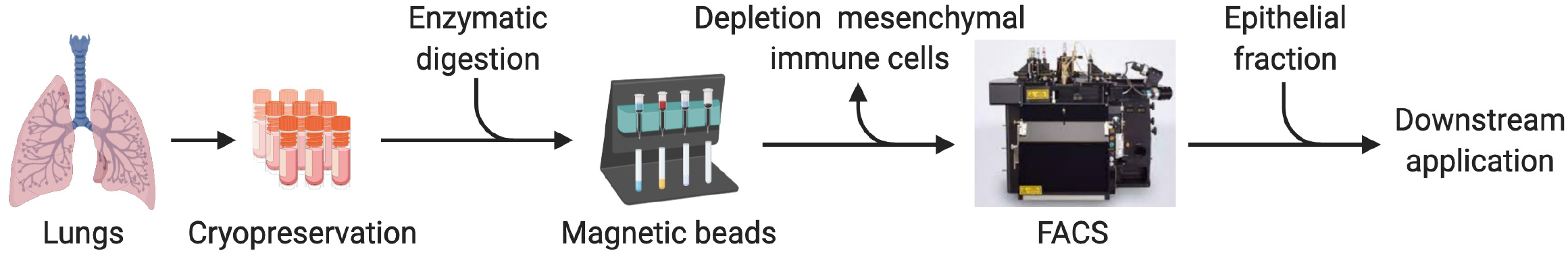
Workflow for isolation of epithelial fraction from human and mouse lung. Cryobanked lung tissue can be used to retrieve single cells by enzymatic dissociation. The epithelial fraction is then obtained by a two steps process involving depletion of immune and endothelial cells with magnetic bead conjugated antibodies, and Fluorescence Activated Cell Sorting with specific epithelial markers.

FACS analysis was performed on the following samples - cells freshly isolated from the mouse lungs (Fig. 2A), cells isolated from Cryobanked minced mouse lung tissue (Fig. 2B), and Cryobanked single cell suspensions of enzymatically dissociated mouse lung tissue (Fig. 2C). For all conditions, epithelial cells were identified based on exclusion of CD31^**+**^ endothelial cells and CD45^+^ immune/hematopoietic cells, followed by positive selection for CD326 (EpCAM). Comparable distributions of CD326+ populations were observed between all samples (Fig. 2A-C; Supp. Fig. 1). Similarly, no significant differences were observed viability of epithelial cells isolated from either fresh (83.53% +/-1.27 %) or Cryobanked (85.00% +/-2.42%) mouse lung tissue (Fig. 2D). However, Cryobanked single cell suspensions of dissociated mouse lung tissue showed a significant reduction in viability compared with either fresh or Cryobanked tissue (p = 0.0008 and 0.0013, respectively). Colony forming efficiency showed no significant differences between epithelial cells isolated from fresh (7.7% +/- 0.9%) or Cryobanked (8.2% +/-1%) tissue but was significantly lower among epithelial cells isolated from Cryobanked single cell suspensions of dissociated mouse lung tissue when compared with either fresh or Cryobanked samples (p = 0.0089 and 0.0007, respectively; Fig. 2E).

**Figure 2:**
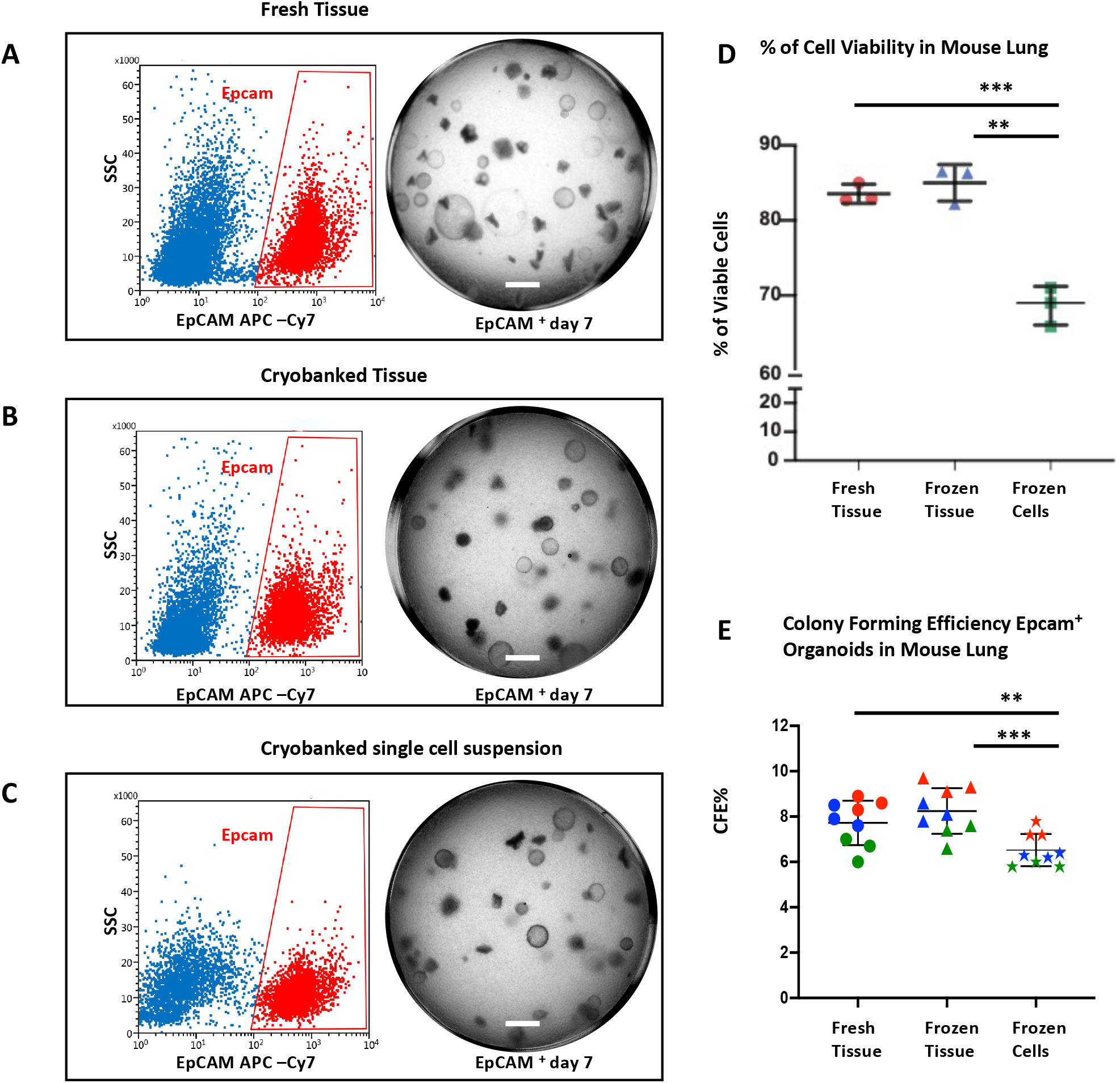
Evaluation of mouse epithelial cell fraction. Representative flow cytometry dot plots of CD31^**-**^/CD45^**-**^/CD326^**+**^ epithelial cells and corresponding 3D organoid cultures from freshly isolated tissue (A), cryobanked tissue (B), or cryobanked single cell suspension (C) of mouse lung (Scale bar = 50 μm). Cell viability (D) P ** = 0.001, P *** = 0.008, and epithelial colony forming efficiency (E) P ** = 0.008, P ***= 0.0007 (n = 3 biological replicates). Error bars represent Standard Deviation and significance determined by unpaired t test.

### Human distal lung cell fractionation comparison fresh tissue Vs frozen tissue Vs frozen cells

Epithelial cells were FACS enriched following enzymatic dissociation of either freshly obtained lung tissue from cadaveric donors (“Distal fresh tissue”, Fig. 3A), or Cryobanked minced tissue (“Distal cryobanked tissue”, Fig. 3B), or from Cryobanked single cell suspensions of enzymatically dissociated tissue (“Distal cryobanked cells”, Fig. 3C). Non-epithelial cell types including immune cells, red blood cells and endothelial cells were depleted from the total pool of lung cells by staining magnetic bead-conjugated antibodies to CD45, CD235a and CD31, respectively. Further FACS depletion of cells staining for either CD45, CD235a and CD31, elimination of dead cells staining for DAPI, and positive selection for the pan-epithelial cell marker CD326, led to highly enriched epithelial cell fraction. Epithelial cells were further subdivided based upon their staining with the AT2-specific monoclonal antibody, HTII-280[41]. Comparable distribution of CD326+ populations were observed between all samples (Fig. 3A-C; Supp. Fig. 2). Similarly, no significant differences were observed viability of epithelial cells isolated from either fresh (86% +/- 6.8%) or frozen (85% +/- 5.8%) human lung tissue (Fig. 3D). HTII 280+ cultures formed rapidly expanding organoids, with an average CFE of 10% in both fresh and frozen distal tissue. Colony-forming efficiency was not significantly different between HTII-280+ cells isolated from fresh (10.7% +/- 2%) and frozen (9.9% +/- 2.8%) human lung tissue (Fig. 3E). In contrast, colony-forming efficiency of HTII-280+ cells isolated from cryobanked single cell suspensions was significantly lower when compared to either fresh (10.7% +/- 2%) or cryobanked tissue (9.9% +/- 2.8% and 7.2% +/- 0.8%, respectively) (Fig. 3E). Distal lung epithelial organoids from fresh tissue (Fig. 3A), frozen tissue (Fig. 3B) and frozen cells (Fig. 3C) were cultured with growth-factor depleted Matrigel in Pneumacult ALI medium. Morphology and immunophenotypic characteristics were equivalent between organoids derived from HTII-280+ cells isolated from either fresh, Cryobanked tissue or Cryobanked single cell suspensions.

**Figure 3:**
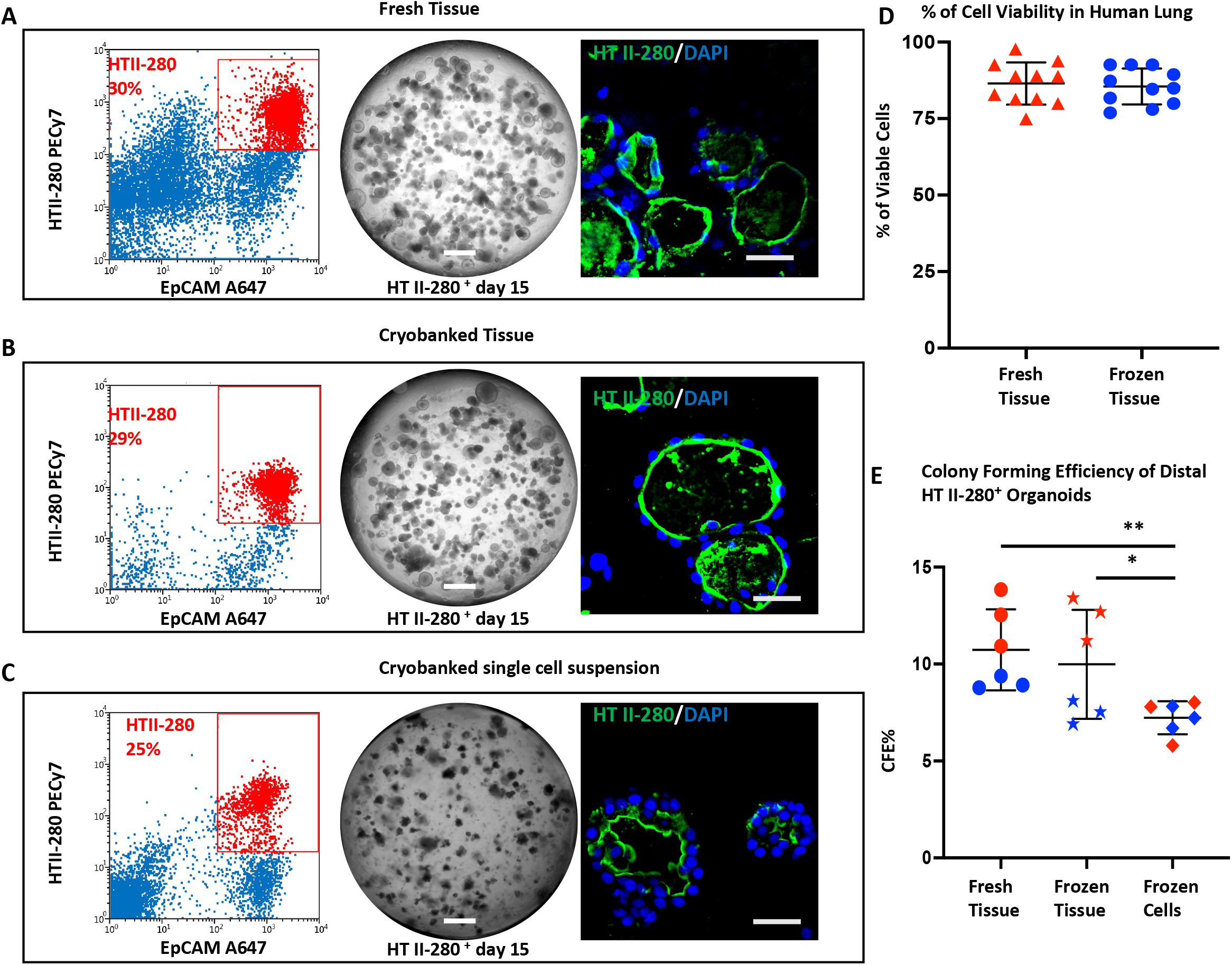
Evaluation of human epithelial cell fraction. Representative flow cytometry dot plots of CD31^**-**^/CD45^**-**^/ CD326^**+**^ /HTII-280^+^epithelial cells and corresponding 3D organoid cultures, and immunoreactivity to HTII-280 are shown for freshly isolated tissue (A), cryobanked tissue (B), or cryobanked single cell suspension (C) of human lung (scale bar = 50 μm). Cell viability (D), and epithelial colony forming efficiency (E) P * = 0.04, P **= 0.003 (n = 3 biological replicates). Error bars represent Standard Deviation and significance determined by unpaired t test.

### Transcriptomic comparison of datasets derived from freshly isolated and frozen tissue

Single cell RNA sequencing was performed to determine the quality of transcriptomic data generated from isolated lung epithelial cells prepared from fresh versus cryobanked lung tissue. Analysis of sequencing quality showed comparable metrics, including sequencing saturation, number of reads mapped to exons, and number of genes per cell (Table 1). Following quality control and filtering, ‘fresh’ and ‘cryobanked’ datasets showed similar representation of transcripts, UMIs, and percentage of mitochondrial genes. Projection of transcript levels for major cell type-specific genes reveal similar cell type-specific profiles among data generated from either fresh or cryobanked tissue samples (Fig. 4B-H). These data suggest that cryobanked tissue samples shown remarkable preservation of transcriptomes of isolated epithelial cell types following recovery and tissue dissociation, relative to their freshly processed tissue counterparts. We conclude that the tissue cryobanking protocol described herein will enable sample banking for simultaneous processing and acquisition of single cell omics data.

**Figure 4:**
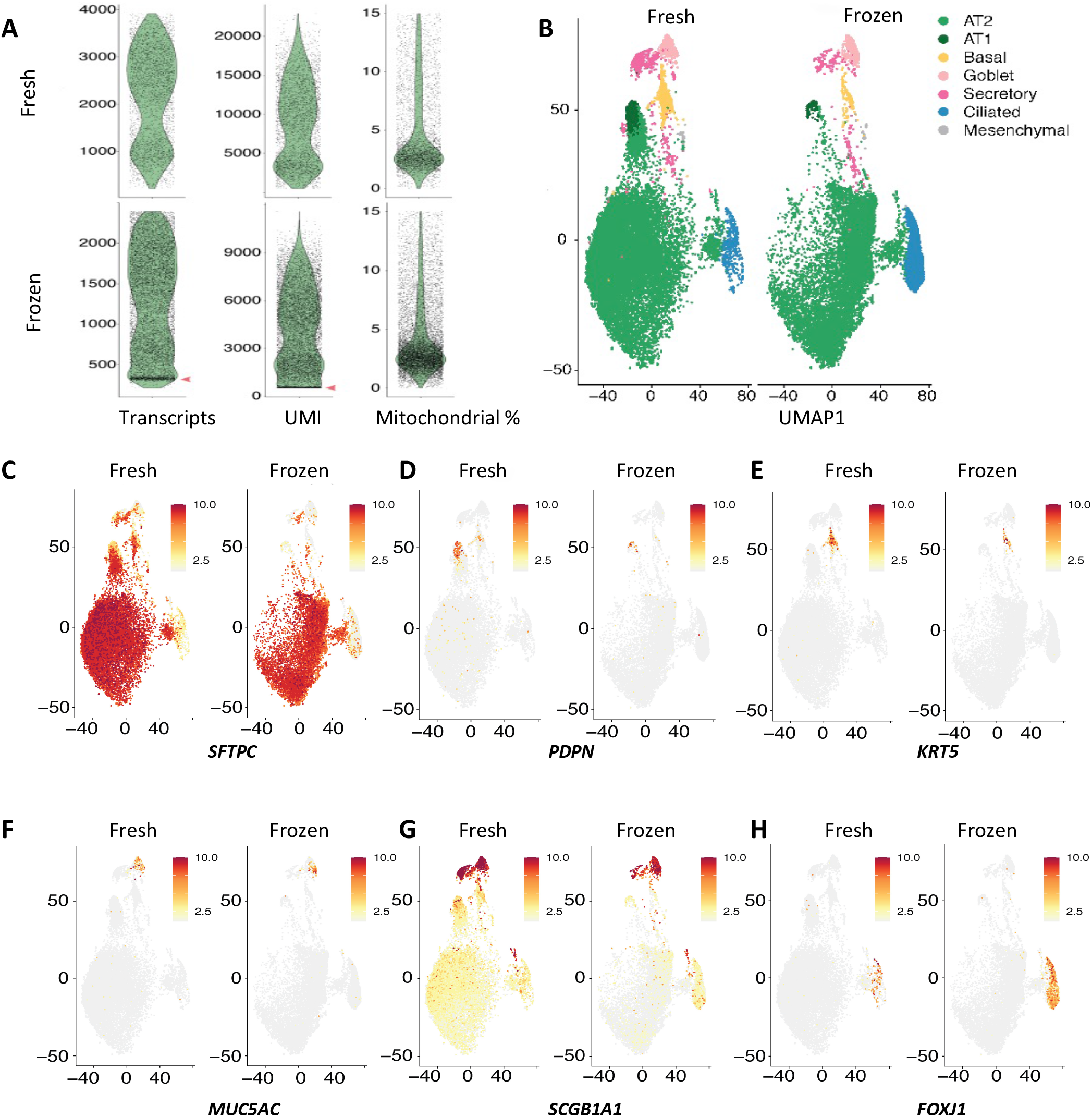
Transcriptomic profiling of fresh and frozen human distal lung tissue. (A) Evaluation of number of transcripts, Unique Molecular Identifiers (UMI), and percentage of mitochondrial genes, in datasets derived from fresh and frozen tissue, visualized by violin plots. Red arrows indicate the presence of a group of cells with low reads for transcripts and UMIs in the ‘frozen’ dataset. (B) Dimensional reduction of data generated from freshly isolated Fresh and Frozen tissue, visualized by UMAP, with cells colored by subset as shown in key. (C-H) Expression of cell-type specific transcripts, divided by ‘fresh’ and ‘frozen’ datasets, visualized by UMAP.

## DISCUSSION

We describe and validate a method for cryopreservation of either mouse or human lung tissue. Epithelial cells isolated from enzymatically dissociated cryobanked tissue retain molecular and functional properties that are equivalent to those of freshly dissociated lung tissue. We have successfully applied this method to tissue from “otherwise healthy” control tissue donors, and explant tissue donated by patients undergoing transplantation for either IPF or CF [8-12, 15] Critical elements of this method include the ease of processing and banking large quantities of tissue, comparable representation of distal lung cell types between cryobanked and fresh tissue samples, preservation of surface epitopes allowing antibody-mediated enrichment of defined cell types, and similar viability of epithelial cells isolated from cryobanked samples to those isolated from fresh tissue.

Epithelial cells recovered from cryobanked tissue maintained the ability to form three-dimensional organoids in culture with an efficiency comparable to epithelial cells isolated from dissociated freshly obtained tissue. Furthermore, single cell RNA sequencing of epithelial cells isolated from cryobanked human lung tissue revealed similar levels of cellular heterogeneity to that seen among epithelial cells isolated from fresh tissue specimens. This held true even for fragile epithelial cell types such as type I pneumocytes and ciliated cells. This is a significant practical advancement as it allows storage of normal and diseased lung tissue based on availability and allows for simultaneous processing of multiple samples for single cell RNA sequencing.

An important benefit of cryobanking multiple patient samples for later simultaneous processing for single cell transcriptomic analysis is the ability to minimize technical variability that contributes to sample batch effects. Comparison of ‘healthy’ and diseased specimens have produced novel description of pathological conditions with unprecedented resolution[8-10]. Collection and processing of large amount of freshly isolated tissue is challenging and often relegated to ad-hoc consortia[16, 17]. In addition, analysis of datasets generated from different institutions, often using laboratory-specific protocols for sample processing, increase the need bioinformatic correction of data to minimize confounding batch effects. While computational method for batch effect correction are available, the ever-growing list of available packages and approaches reflect that technical challenges associated with their application, the principal caveat being that batch effect removal is frequently associated with data degradation and loss of significant biological differences. Cryopreservation has been shown in other organ systems to allow serial sample collection with simultaneous tissue processing to minimize technical batch effects [18]. Here we developed and validated a method that is specific for the lung, including distal epithelium, allowing sample collection, and sharing between remote laboratories for simultaneous processing to guarantee uniformity for omics analyses.

In summary, newly developed methods described herein allow standardization of tissue procurement and experimental planning that are important requirements for development of uniform culture methodologies, screening assays, as well as for assessment of potential applications for lung regenerative medicine. These methods provide a tool for large-scale banking of valuable lung tissue samples for sharing within the research community.

## Acknowledgments

We acknowledge support from the Applied Genomics, AHSP/RMI Flow Cytometry, and Biobank cores at Cedars Sinai Medical Center (CSMC). This research was supported by grants from the National Institutes of Health (NIH) (R01 HL135163; P01 HL108793), the Cystic Fibrosis Foundation (STRIPP20XXO, CARRAR19G0), the Bristol Myers Squibb IDEAL consortium, and by the Bram and Elain Goldsmith Chair in Gene Therapeutics Research.

## Author contributions

Conceptualization, B.K, G.C, B.R.S.; methodology, B.K., G.C., C.Y., E. I., B.R.S.; tissue procurement, C.A.H., W.G.T., P.C., R.R., S.B.; data analysis B.K, G.C, B.R.S.; writing, B.K, G.C, B.R.S.; funding acquisition, B.R.S.

## Supplemental data

**Table 1:**
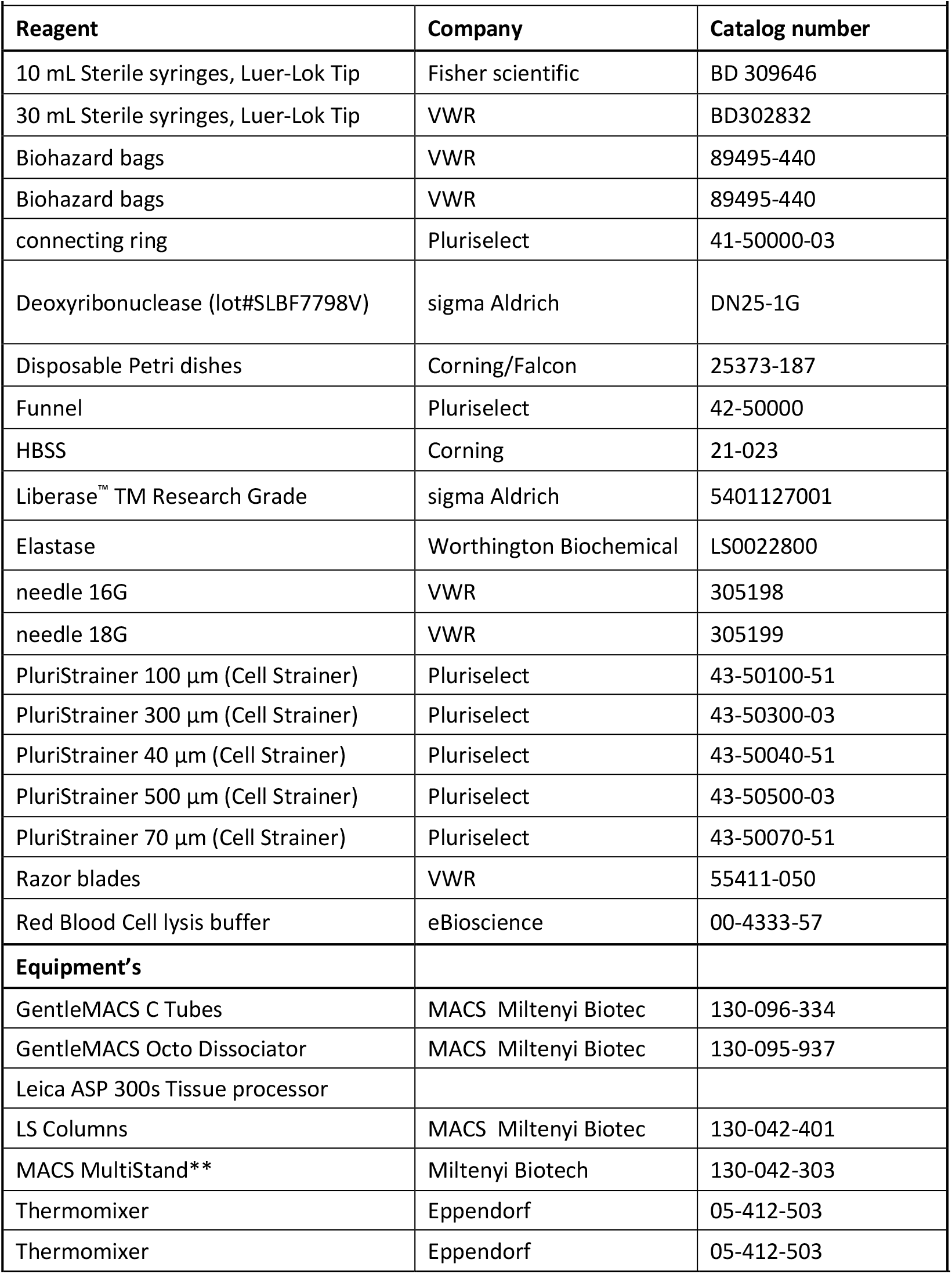
Cell Isolation.

**Table 2:**
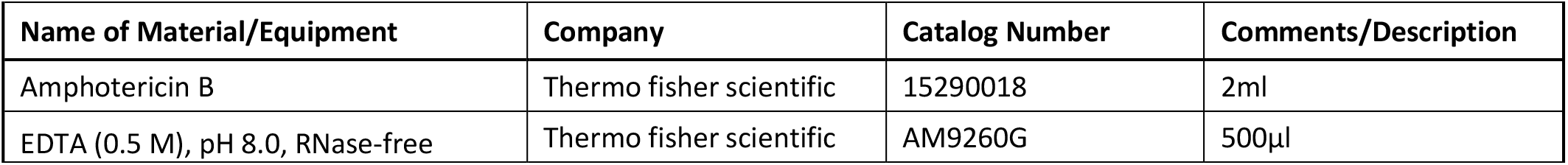
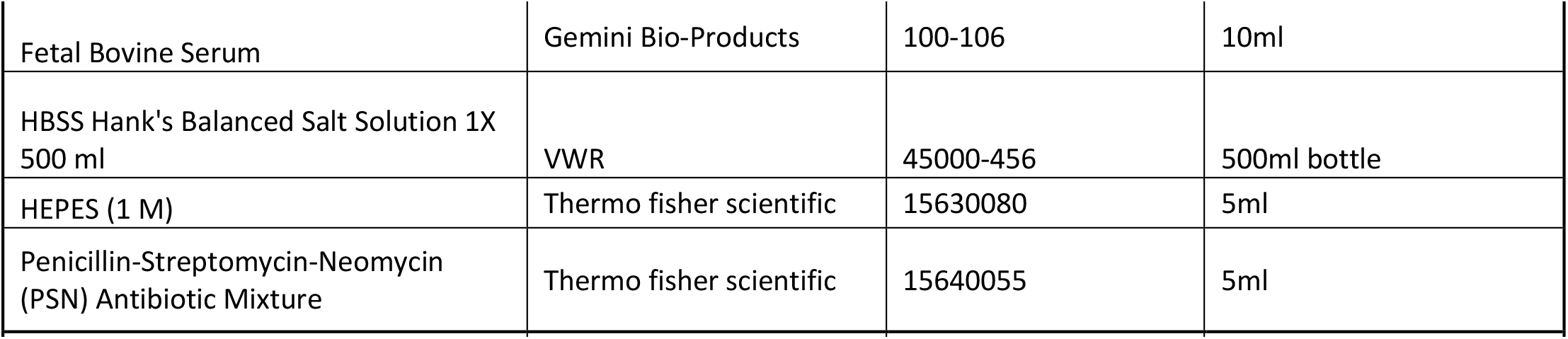
FACS Buffer.

**Table 3:**
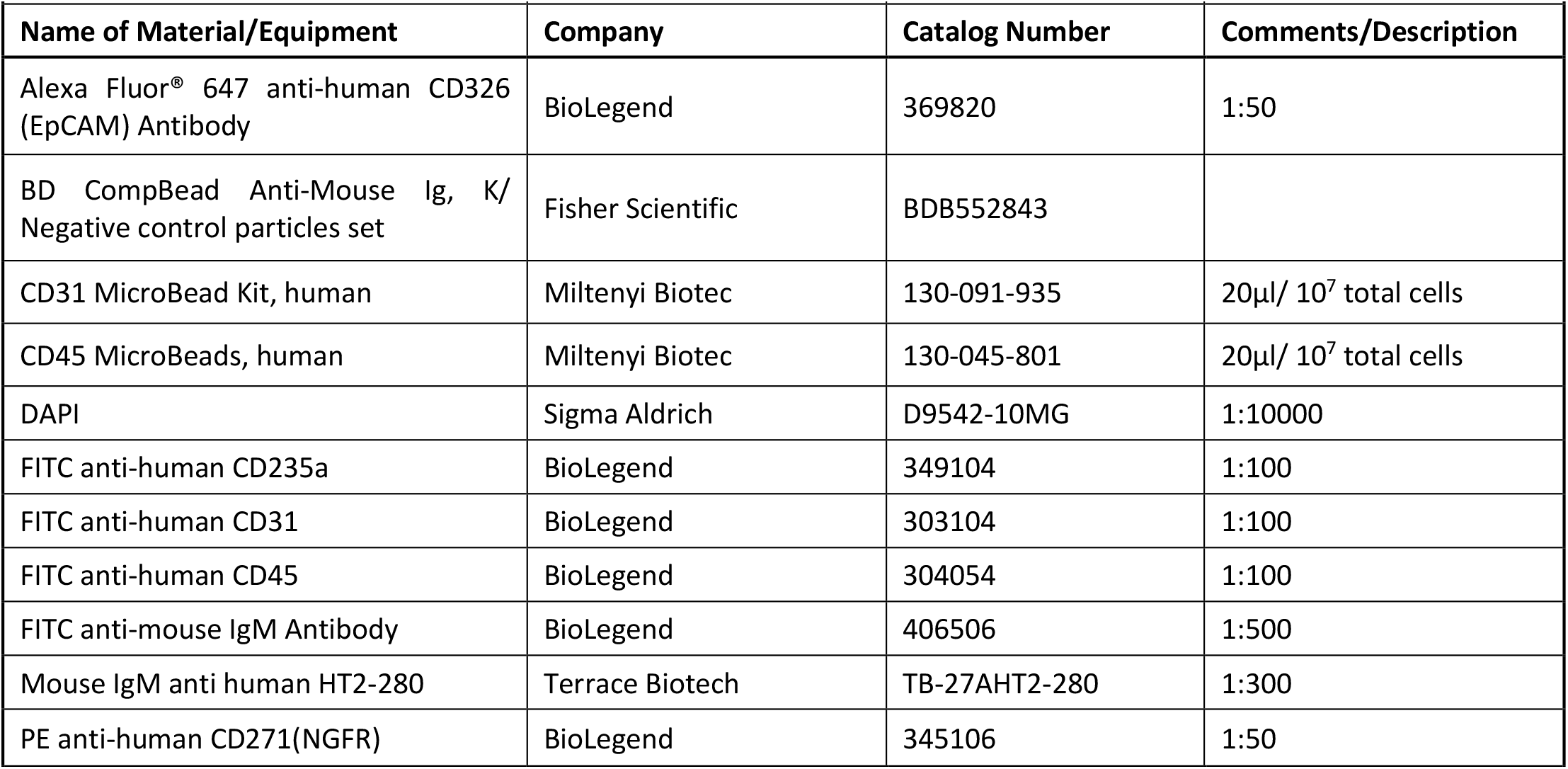
List of antibodies for FACS.

**Table 4:**
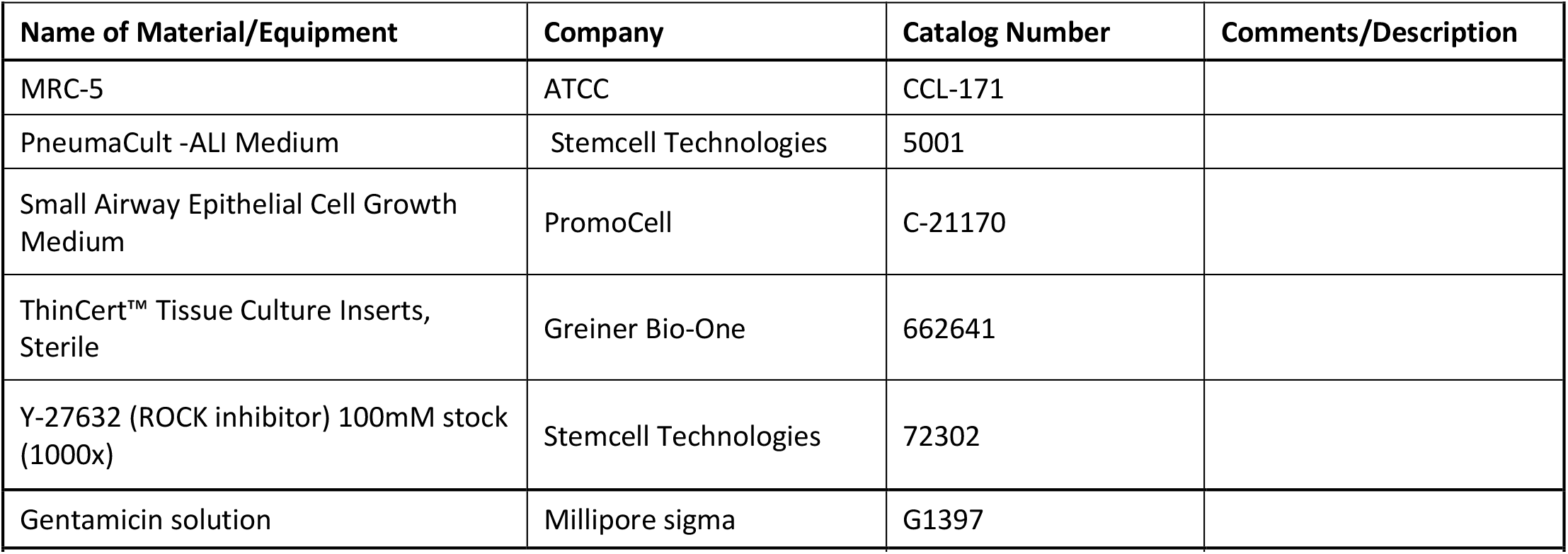
Composition of Organoid Culture mediums.

**Table 5:**
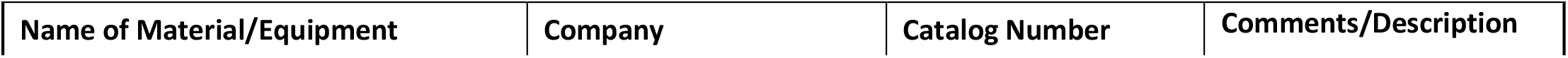
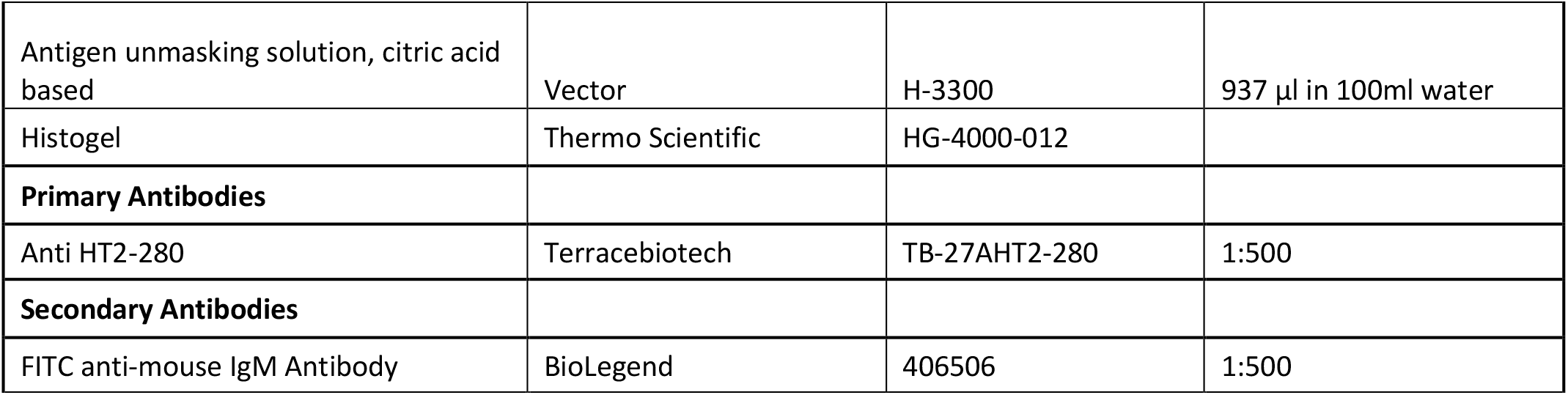
List of antibodies for Immunohistochemistry.

**Table 6:**
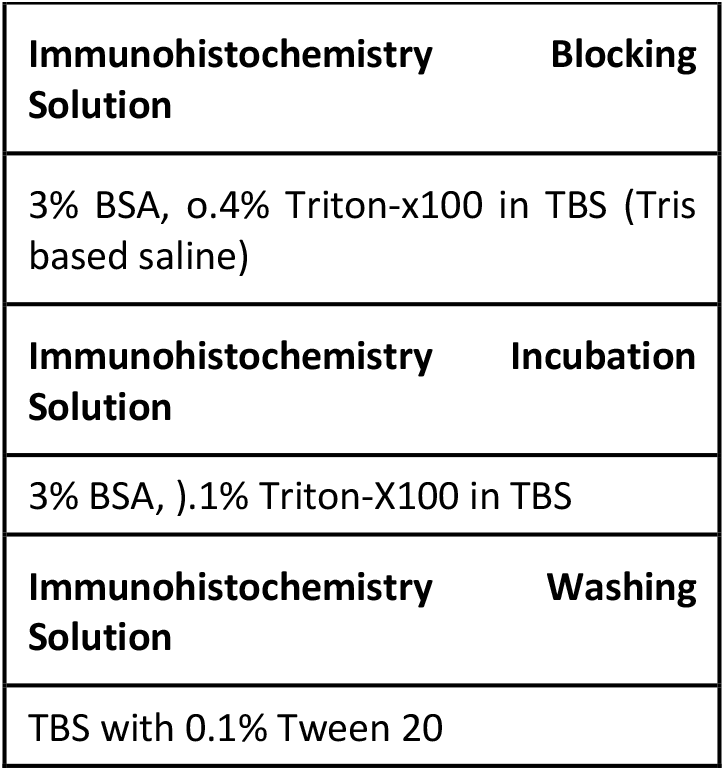
Buffers.

## Supplemental Data

### Mouse Lung

#### Isolation of cells from Mouse Lung

1. **Mouse lung isolation**. Mice were placed in a surgical plane by I.P. injection of Ketamine (100 mg/kg) and Xylazine (10 mg/kg) and exsanguinated by dissection of the aorta. Lungs were deflated by nicking the diaphragm and a catheter (20G) inserted into the trachea. Lungs were lavaged three times with 1 ml ice-cold PBS. Lung tissue was then removed from the thorax and immersed in ice cold PBS. Mouse lung epithelial cells were isolated as previously described, and summarized below. [42].
2. **Mouse Lung tissue dissociation**. Lung tissue was suspended in PBS at 37 °C and inflated with Hams F12 medium containing elastase (4 U/ml). Additional Hams F12/Elastase was introduced every 5 minutes for a total incubation time of 20 minutes. Lung tissue was minced in a petri dish to yield lung fragments of <1mm, and further enzymatically digested by the addition of Liberase TM (0.25 Wünsch U/ml) in HBSS at 37 °C for 60 min. The digested tissue was then passed through a 70 μm strainer, resuspended in HBSS-FACS buffer and centrifuged at 400 g for 5 mins at 4ºC. Next the pellet was resuspended in 1ml of Red Blood Cell (RBC) lysis buffer and incubated for 1.5 mins with gentle agitation at RT, and then passed through a 70 μm strainer. Finally, cells were centrifuged at 400 g for 5 mins at 4°C and resuspended in HBSS-FACS buffer.
3. **Depletion of immune and endothelial populations**. Microbeads for immune (CD45) and endothelial (CD31) cell removal (10 μl each), were added to 90 μl of HBSS-FACS buffer containing 10^7^ total cells. After a 15-20 mins incubation at 4 °C, cells were washed in HBSS-FACS buffer and centrifuged at 300×g for 5 mins. A LS Column, placed in the magnetic field of a MACS Separator, was primed by rinsing with 3 mL of HBSS-FACS buffer, then the cell suspension was added onto the column, and effluent unlabeled cells were collected and used for FACS.
4. **Cell Surface staining for FACS**. Per each 1 × 10^7^ cells in 1 mL of HBSS+ buffer, primary antibodies were added at the required concentration and incubated for 30 mins at 4 °C. (See antibody details in table 3). Next, cell were washed in HBSS+ buffer and centrifuged at 600 x g for 5 mins at 4°C. Finally, before proceeding to FACS, cells were filtered through a strainer to ensure single cell suspension, and DAPI (1μg/mL) was added to stain for dead cells.

## Supplemental Figures

**Supplemental Figure 1:**
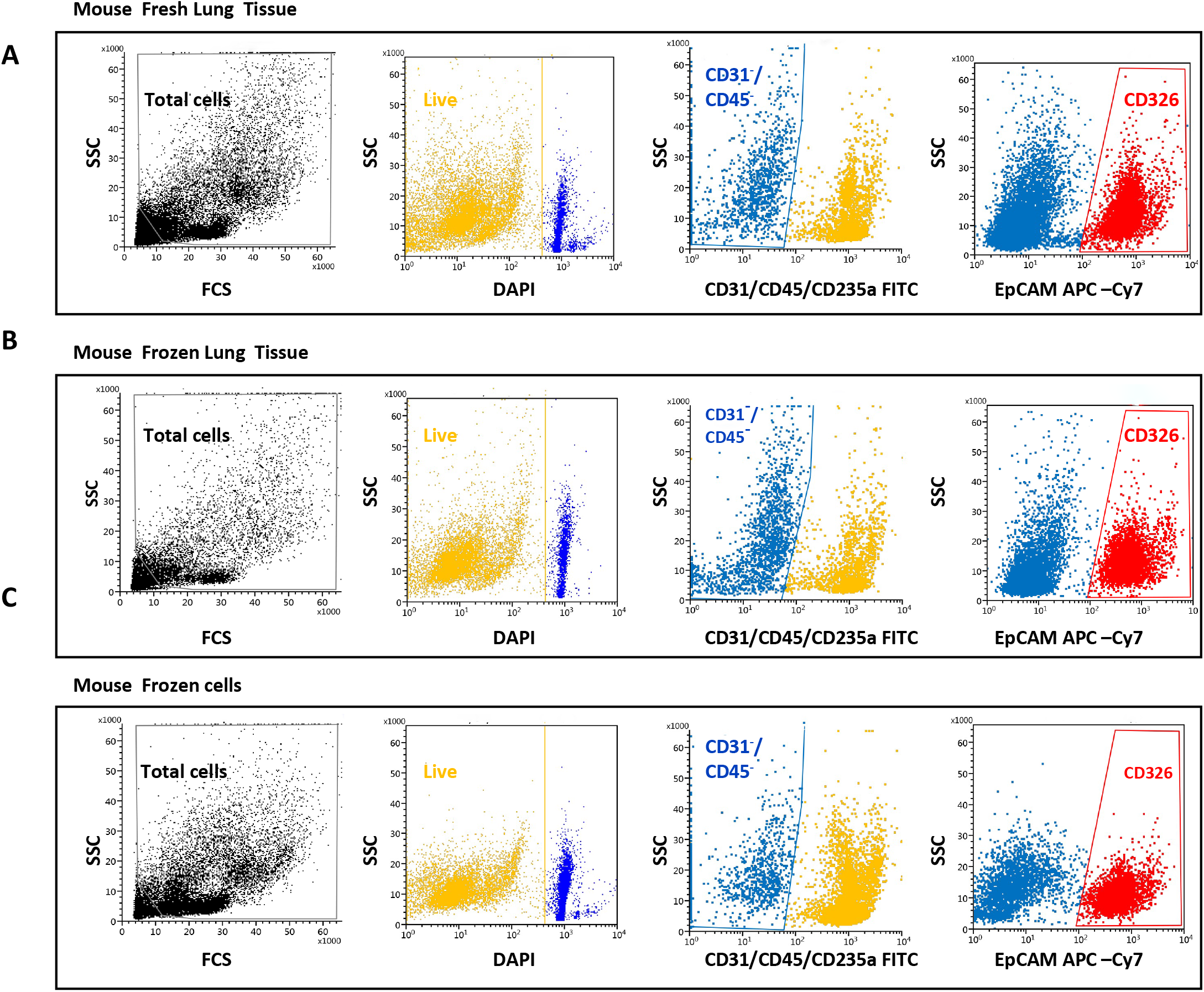
Gating strategy for mouse lung epithelial cell fractionation. Representative image showing the comparison between FACS plots of CD31^**-**^/CD45^**-**^/CD326^**+**^ population from fresh tissue (A), cryobanked frozen tissue (B) and cryobanked single cell suspension (C) from a single mouse lung.

**Supplemental Figure 2:**
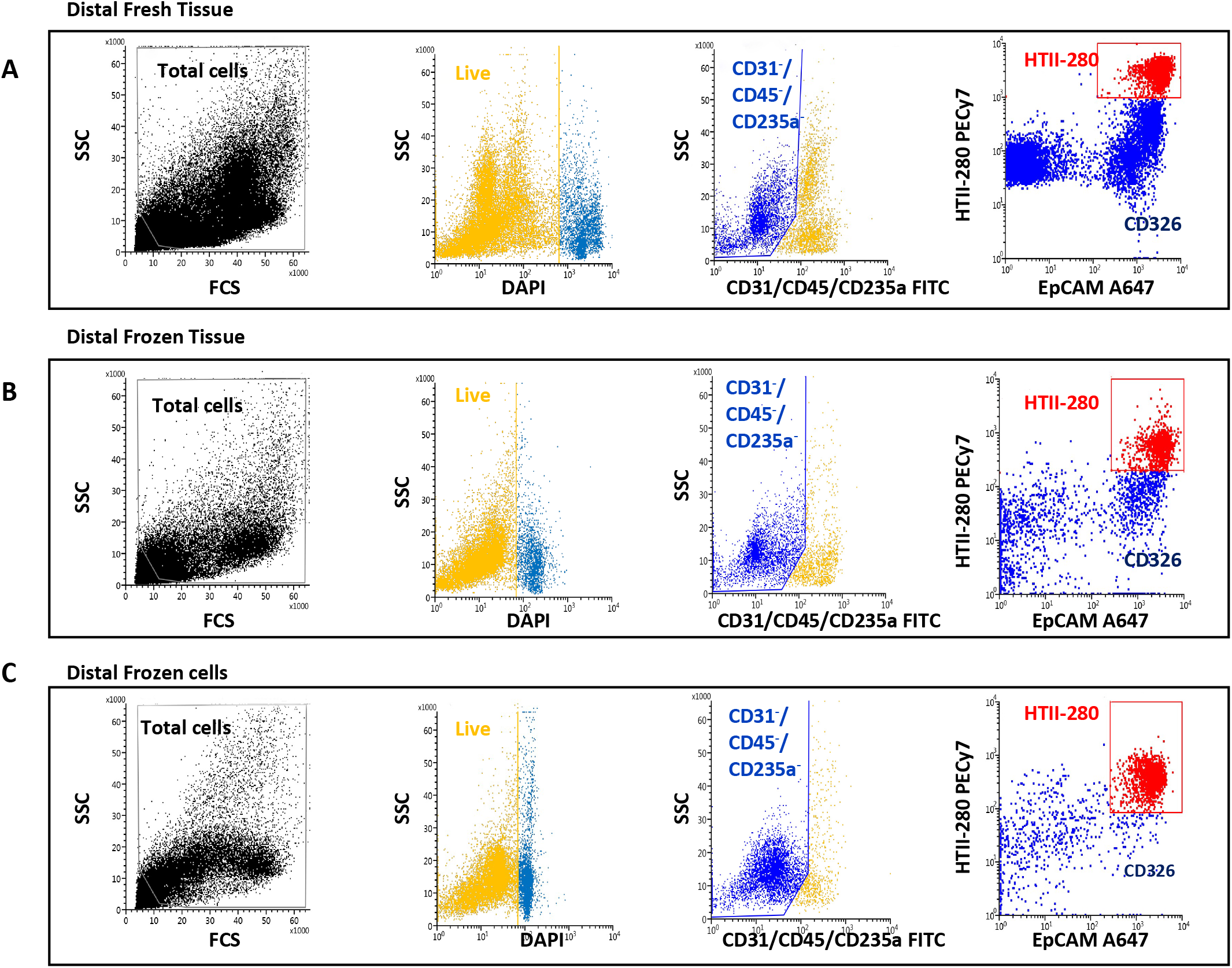
Gating strategy for human lung epithelial cell fractionation: Representative image showing the comparison between FACS plots of CD31^**-**^/CD45^**-**^/CD326^**+**^**/**HTII-280^**+**^ population from fresh distal tissue (A), cryobanked frozen distal tissue (B) and cryobanked single cell suspension from distal tissue(C) from a single biological sample of human distal lung.

**Supplemental Figure 3:**
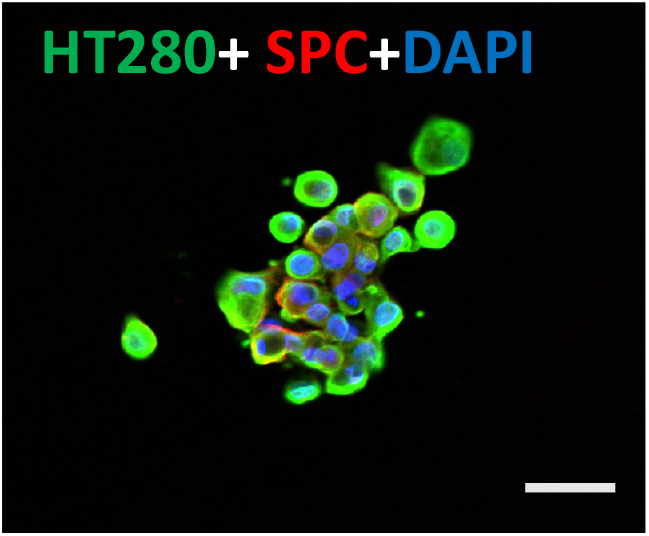
Sorted cells were stained for HTII-280 (green), SPC (red) and nuclei (blue). Scale bar = 50 μm.

